# Effect of pesticides and metals on zebrafish embryo development and larval locomotor activity

**DOI:** 10.1101/2020.10.05.326066

**Authors:** Farooq Ahmad, Shaukat Ali, Michael K. Richardson

**Affiliations:** Institute of Biology Leiden (IBL), Leiden University Sylvius Laboratory, Sylviusweg 72, 2333 BE, Leiden, The Netherlands; Present address: Department of Zoology, The Islamia University of Bahawalpur, Pakistan; Present address: Department of Zoology, Government College University, Lahore-54000, Lahore, Pakistan

## Abstract

The zebrafish has been widely used as a predictive model in safety and toxicology. Low cost high-throughput screening can be achieved with this model, and the genome contains orthologues of the majority of human disease genes. However, previous studies indicate that the predictivity of the zebrafish model in toxicology varies between compound and compound class. We examined this issue by screening 24 compounds from two different compound classes, metals and biocides (pesticides/insecticides) for toxicity in the zebrafish model and looked at the effects on hatching, morphology and predictivity for mammalian toxicity. Wild-type zebrafish embryos were exposed to test compounds in 96-well plates for 96 hours starting at 24 hours post fertilization. Hatching was either delayed or accelerated depending on the compound. Three types of alteration in behavioural responses were noted: (i) hypoactivity; (ii) hyperactivity; and (iii) biphasic response (a dose-dependent shift between hypo- and hyperactivity). LC_50_ of compounds was calculated and compared to published LD_50_ values in rodents. The zebrafish-rodent values were poorly correlated for both metals and biocides. We conclude that, although the zebrafish is a good model for some aspects of toxicology, its predictivity for mammalian toxicity needs to be determined per compound class.

## Introduction

The zebrafish is a small, teleost fish of shallow, fresh-water, which has emerged as a valuable model in the field of research especially in the last decade (1). The advantages which have made it a popular model in research are manifold and include: external fertilisation and rapid development, low maintenance costs, easy, year-round spawning, rapid generation cycle (2–3 months), and ease of use for high-throughput screening (2). Its genome is also nearly completely sequenced and contains orthologues of 82% of human disease genes (3, 4). The zebrafish is used in many fields of biology research including behaviour (5–8), chemical toxicity (9–14), drug discovery (15–17) and in human disease modelling (18–21) by using forward and reverse genetic techniques together with large-scale, high-throughput screening. However, more information is needed on the predictivity of the zebrafish model in toxicity, that is, to what extent does the toxicity of compounds tested on zebrafish correlate with their toxicity in mammals (especially rodents and humans)?

Given the aforementioned advantages of the zebrafish, the effects of both short- and long-term exposure to a wide range of toxins can be studied with relative ease. A variety of compounds has been tested on zebrafish, and includes metals and organic compounds (22, 23) and mixtures of drugs (24). The main emphasis in these studies has been on lethality, embryo survival rate and organ malformation as general assay parameters, and demonstrated that zebrafish exhibit good dose-responsiveness to toxicity and are a suitable animal model for toxicity screening (14, 25, 26).

The use of zebrafish in behavioural neuroscience is in its infancy compared to the use of rodents (27). However, mutant zebrafish lines, morpholinos, high-throughput screening and new bioassays for toxic and therapeutic endpoints in zebrafish are likely to become more common. New technology is having a large impact on research, and this will result in greater insights into the mechanisms of toxicity of chemicals, as well as aiding in the discovery of new drugs for treating several human diseases (27–29). Although the number of published studies on zebrafish behaviour is not large compared to comparable studies on rodents, many of the behaviours displayed by zebrafish are well-described. These include the open-field test (30, 31), optomotor response (32), optokinetic response (33–37), photokinesis (5) and visual motor response test (38–40) among many others.

It has long been known that behavioural patterns of animals including zebrafish can be altered by drugs and chemicals (41–43). These alterations are regarded as an observable expression of effects on nervous and locomotor systems (13). Some of the environmental chemicals, such as pesticides, can cause developmental neurotoxicity resulting in neurodevelopmental disorders in humans (44, 45). This makes it important to determine the effects of these chemicals on living animals and their behaviour.

Several classes of compound have been tested on zebrafish and assessed for their toxicity prediction in rodents. The predicitivity was found to vary considerably according to compound or compound class (9, 46, 47). In the current study, we have tested metals, pesticides and insecticides (and the latter two we shall collectively call ‘biocides’) on zebrafish embryos. We have compared the results with studies of toxicity of the same compounds in mammals. We chose these compounds because they are very diverse chemically and because there is increasing awareness and concern regarding the environmental effects of these compounds (48, 49). For these reasons, the predictivity of the zebrafish in relation to the toxicity of these compounds in mammals is an important consideration, because it is a potential test model in environmental toxicology.

## Material and methods

### Statement of ethics on animal use

All experimental procedures were conducted in accordance with The Netherlands Experiments on Animals Act that serves as the implementation of "Guidelines on the protection of experimental animals" by the Council of Europe (1986), Directive 86/609/EC, and were performed only after a positive recommendation of the Animal Experiments Committee had been issued to the license holder.

### Animal husbandry

Wild-type male and female adult zebrafish (*Danio rerio*) were purchased from Selecta Aquarium Speciaalzaak (Leiden, The Netherlands) who obtains stock from Europet Bernina International BV (Gemert-Bakel, The Netherlands). We limited our experiment to AB strain of zebrafish as different strains have differences in the locomotor activity (50). Fish were kept at a maximum density of 12 individuals in plastic 7.5 L tanks (1145, Tecniplast, Germany) containing a plastic plant as tank enrichment, in a zebrafish recirculation system (Fleuren & Nooijen, Nederweert, The Netherlands) on a 14h light: 10h dark cycle (lights on at 7h AM: lights off at 21h PM). Water and air temperature were maintained at 24 ^°^C and 23 ^°^C, respectively. Fish were purchased at the juvenile stage and were allowed to adapt to our facility for at least 2 months before being used as adult breeders. The fish were fed daily with dry food (DuplaRin M, Gelsdorf, Germany) and frozen artemias (Dutch Select Food, Aquadistri BV, The Netherlands).

Zebrafish eggs were obtained by random mating between sexually mature individuals. Briefly, on the day (16h) before eggs were required, a meshed net allowing eggs to pass through but preventing adult fish from accessing/eating them, was introduced in the home tank of a group of 12 adult fish. Each breeding tank was only used once per month to avoid handling stress and ensure optimal eggs quantity and quality.

The eggs were harvested the next day (30 min after the onset of lights at 7h AM) and age was set as post fertilization day (dpf) 1 based on the staging system employed in the zebrafish text book entitled *Zebrafish: a practical approach* (51). They were placed in 9.2 Petri dish containing 100 ml egg water (0,21 g/l Instant Ocean Sea Salt and 0,0005% (v/v) methyl blue). 50-60 eggs were place in one Petri dish in a climate room maintained at a temperature of 28 °C and 50% humidity and under a light-dark cycle of 14h:10h (lights on at 7h AM/lights off at 9h PM).

### Zebrafish Egg plating

At 24 hours post fertilization (hpf), all embryos were checked for their natural spontaneous mortality as there are reports of an early natural death in zebrafish embryos cultured under certain conditions (9, 10). In order to avoid taking embryos during such a die-off, we used 24h old embryos for exposure of chemicals after removing unfertilized eggs and refreshing the egg water. Thus, each larva was gently taken up into a plastic Pasteur pipette (VWR International B.V., The Netherlands) and directly transferred to 96-well plate, one larva per well containing 250 µl egg water (control) or respective concentration of compound tested. Note that in order to eliminate further sources of disturbance/stress, the media was not refreshed except on 2dpf where the medium was completely replaced by fresh egg water and non-fertilized eggs were removed. At the end of the behavioural testing, the larvae were processed further as follows for morphological assessment.

### Morphological assessment

Embryos were fixed in 4% paraformaldehyde (PFA) in phosphate-buffered saline at pH 7.2 at 4°C overnight. They were then rinsed five times in distilled water and dehydrated in a graded series of ethanol (25, 50, and 70%) for 5 min each. Embryos were rinsed in acid alcohol (1% concentrated hydrochloric acid in 70% ethanol) for 10 min. They were then placed in filtered Alcian blue solution (0.03% Alcian blue in acid alcohol) overnight. Embryos were subsequently differentiated in acid alcohol for 1 h and washed 2×30 min in distilled water. All embryos remained in their original multiwall plates, so that each individual could be tracked throughout the entire experimental and analysis procedure. Analysis of embryo morphology was carried out using a dissecting stereo microscope. The phenotypes of malformations scored are defined in Table 1.

**Table 1:** Phenotypic endpoints scored in embryos at 5dpf. Some of these criteria have been described elsewhere (52).

| Phenotype | Criteria |
| --- | --- |
| Yolksac | Yolksac enlarged by fluid accumulation |
| Heart | Pericardial sac enlarged by fluid accumulation |
| Meckel cartilage | Meckel cartilage grossly hypoplastic, missing or unfused in midline. |
| Melanocytes | Melanocytes aggregate and dispersed |
| Short body | Body was considered short if its total length was below 3.5mm at 6dpf |
| Tail | A bent on tail. |
| Body axis | The trunk was bent on dorso-ventral axis |
| Normal | The larvae was considered with normal morphology if any of the above listed phenotypes were normal |

**Table 2:**
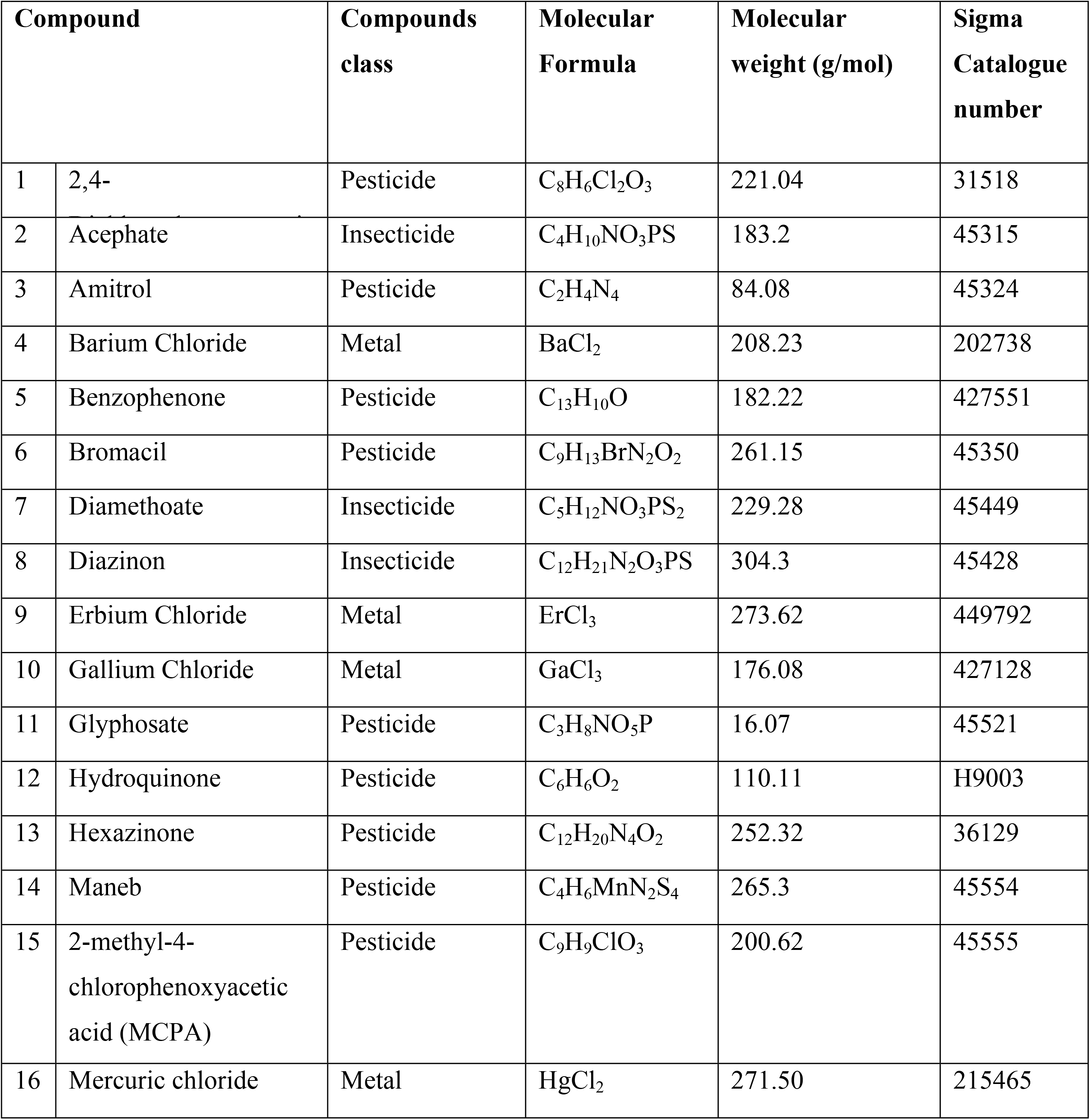
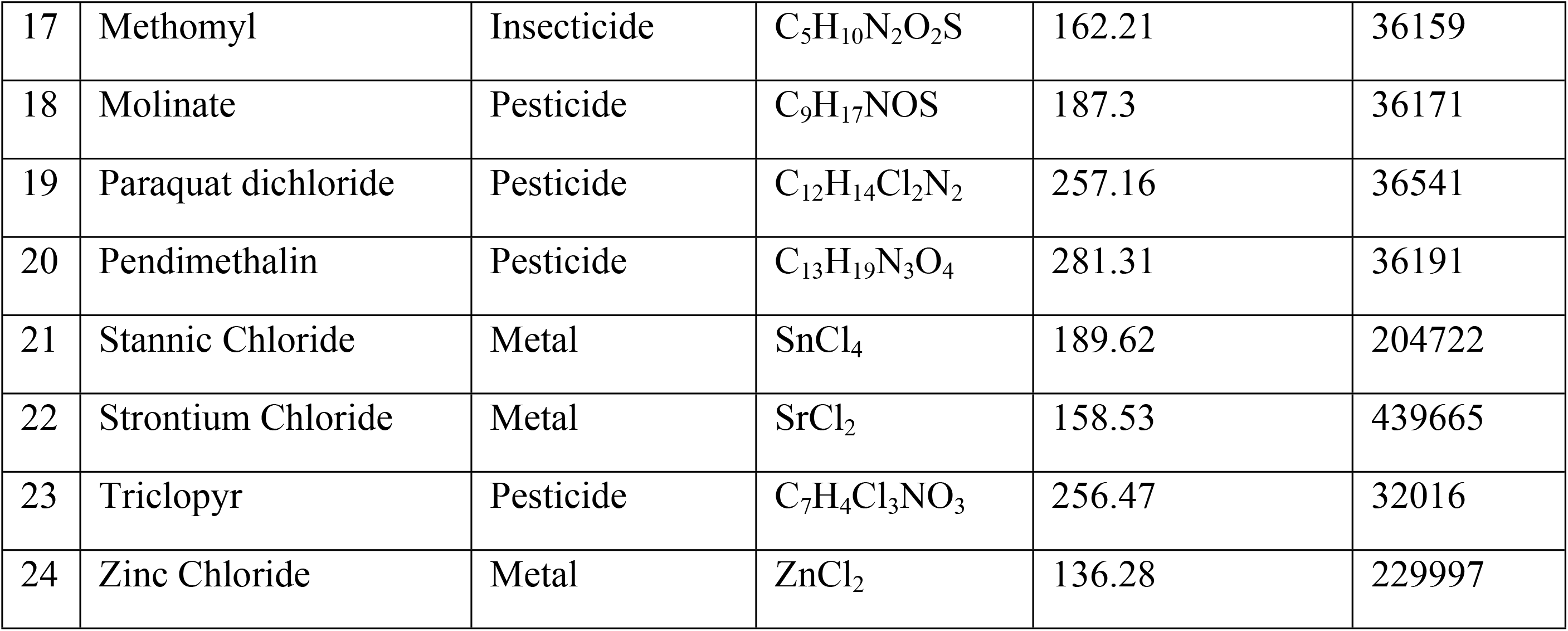
List of compounds used in the study; all compounds were purchased from Sigma (Zwijndrecht, The Nethelrands).

| Compound | Compounds class | Molecular Formula | Molecular weight (g/mol) | Sigma Catalogue number |
| --- | --- | --- | --- | --- |
| 1 2,4- | Pesticide | C<sub>8</sub>H<sub>6</sub>Cl<sub>2</sub>O<sub>3</sub> | 221.04 | 31518 |
| 2 Acephate | Insecticide | C<sub>4</sub>H<sub>10</sub>NO<sub>3</sub>PS | 183.2 | 45315 |
| 3 Amitrol | Pesticide | C<sub>2</sub>H<sub>4</sub>N<sub>4</sub> | 84.08 | 45324 |
| 4 Barium Chloride | Metal | BaCl<sub>2</sub> | 208.23 | 202738 |
| 5 Benzophenone | Pesticide | C<sub>13</sub>H<sub>10</sub>O | 182.22 | 427551 |
| 6 Bromacil | Pesticide | C<sub>9</sub>H<sub>13</sub>BrN<sub>2</sub>O<sub>2</sub> | 261.15 | 45350 |
| 7 Diamethoate | Insecticide | C<sub>5</sub>H<sub>12</sub>NO<sub>3</sub>PS<sub>2</sub> | 229.28 | 45449 |
| 8 Diazinon | Insecticide | C<sub>12</sub>H<sub>21</sub>N<sub>2</sub>O<sub>3</sub>PS | 304.3 | 45428 |
| 9 Erbium Chloride | Metal | ErCl<sub>3</sub> | 273.62 | 449792 |
| 10 Gallium Chloride | Metal | GaCl<sub>3</sub> | 176.08 | 427128 |
| 11 Glyphosate | Pesticide | C<sub>3</sub>H<sub>8</sub>NO<sub>5</sub>P | 16.07 | 45521 |
| 12 Hydroquinone | Pesticide | C<sub>6</sub>H<sub>6</sub>O<sub>2</sub> | 110.11 | H9003 |
| 13 Hexazinone | Pesticide | C<sub>12</sub>H<sub>20</sub>N<sub>4</sub>O<sub>2</sub> | 252.32 | 36129 |
| 14 Maneb | Pesticide | C<sub>4</sub>H<sub>6</sub>MnN<sub>2</sub>S<sub>4</sub> | 265.3 | 45554 |
| 15 2-methyl-4-chlorophenoxyacetic acid (MCPA) | Pesticide | C<sub>9</sub>H<sub>9</sub>ClO<sub>3</sub> | 200.62 | 45555 |
| 16 Mercuric chloride | Metal | HgCl<sub>2</sub> | 271.50 | 215465 |

|  |  |  |  |  |  |
| --- | --- | --- | --- | --- | --- |
| 17 | Methomyl | Insecticide | C<sub>5</sub>H<sub>10</sub>N<sub>2</sub>O<sub>2</sub>S | 162.21 | 36159 |
| 18 | Molinate | Pesticide | C<sub>9</sub>H<sub>17</sub>NOS | 187.3 | 36171 |
| 19 | Paraquat dichloride | Pesticide | C<sub>12</sub>H<sub>14</sub>Cl<sub>2</sub>N<sub>2</sub> | 257.16 | 36541 |
| 20 | Pendimethalin | Pesticide | C<sub>13</sub>H<sub>19</sub>N<sub>3</sub>O<sub>4</sub> | 281.31 | 36191 |
| 21 | Stannic Chloride | Metal | SnCl<sub>4</sub> | 189.62 | 204722 |
| 22 | Strontium Chloride | Metal | SrCl<sub>2</sub> | 158.53 | 439665 |
| 23 | Triclopyr | Pesticide | C<sub>7</sub>H<sub>4</sub>Cl<sub>3</sub>NO<sub>3</sub> | 256.47 | 32016 |
| 24 | Zinc Chloride | Metal | ZnCl<sub>2</sub> | 136.28 | 229997 |

### Egg water

Egg water was made from 0.21 g ‘Instant Ocean^®^’ salt in 1 L of Milli-Q water with resistivity of 18.2 MΩ cm.

### Test COMPOUNDS

The compounds used in the present study are listed in the **Error! Reference source not found.**.

### Range-finding test

A range-finding test was conducted using a logarithmic series to determine a suitable range of concentration (0, 1, 10, 100 and 1000 mg/L) as recommended in standard protocols (53). After 24hpf, zebrafish embryos were checked for the natural mortality and after removing dead embryos, healthy ones were transferred from Petri dish using a sterile plastic pipette into 96-well microtitre plates. A single embryo was placed in each well so that dead embryos would not affect others, and also to allow individual embryos to be tracked for the whole duration of the experiment. We used a static non-replacement regime without any replacement or refreshment of egg water or test compound. Each well contained 250 mL of either freshly prepared test compound; egg water (control) or vehicle (egg water with solvent where mentioned). 16 embryos for each concentration and 16 embryos as controls for each compound were used.

### Geometric series and LC_50_ determination

A geometric series was selected based on the mortality rate of the range-finding series with concentrations lying in the range 0-100% mortality. The actual concentrations used are shown inxs Table S1. The concentrations were in a geometric series in which each was 50% greater than the next lowest value as recommended (53). Each compound was tested in triplicate (48 embryos per concentration and 48 embryos for control and/or vehicle for each compound). LC_50_ (expressed in mg/L of egg water) was determined based on cumulative mortality obtained from three independent experiments at 120 hpf using Regression Probit analysis with SPSS Statistics for windows version 17.0 (SPSS Inc., Chicago, USA). The embryos were exposed to the compound for 96 h as in the range finding test. The LC_50_ in mg/L was converted into LC_50_ mmol/L to make relative toxicity easier to examine.

### Hatching and Mortality scoring

Hatching was monitored from 48-72 hpf which is the normal hatching period of zebrafish larvae (54). The hatching rate was recorded once all the embryos in any particular concentration were hatched.

Mortality rate (Table 3) was recorded at 48, 72, 96 and 120 hpf in both logarithmic series and geometric series using a dissecting stereomicroscope. Embryos were scored as ‘dead’ if there was no locomotor activity, the heart stopped beating and the change in appearance of tissues from a transparent to opaque.

**Table 3:**
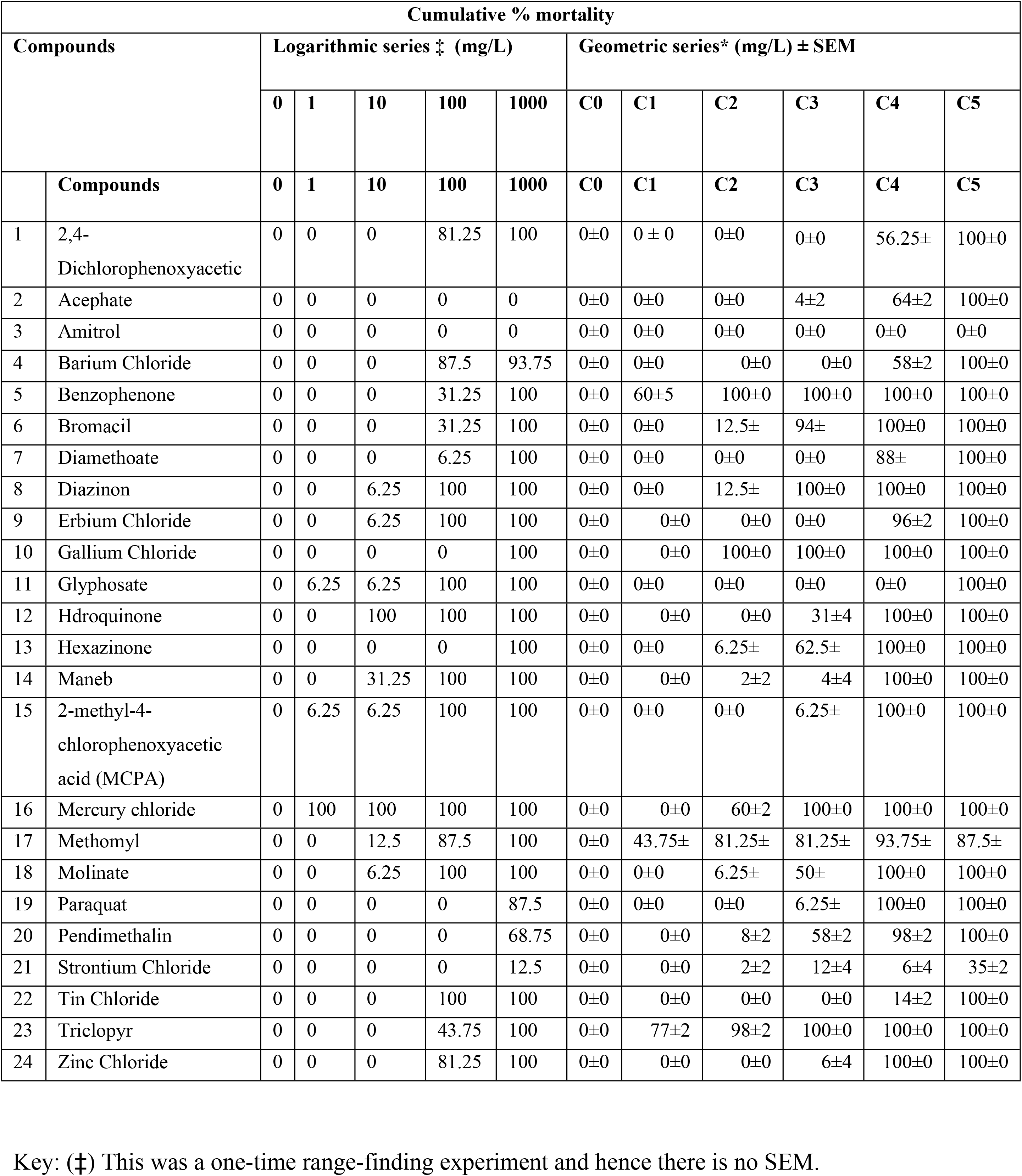
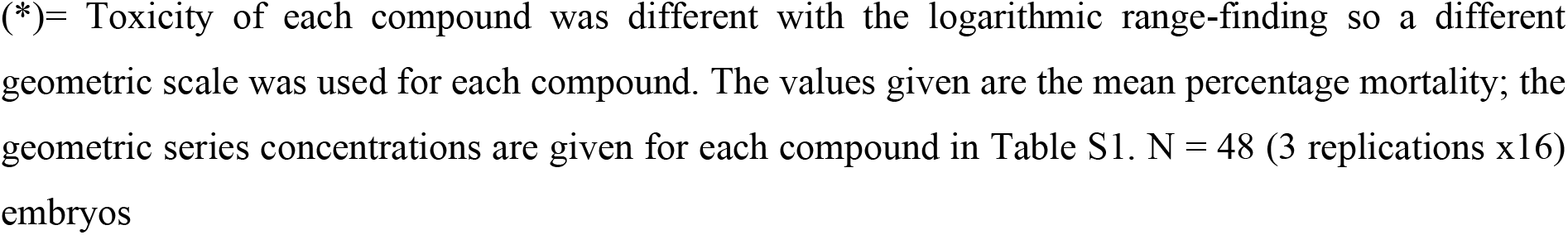
Cumulative % mortality recorded in 5d larvae after 96 h exposure

### Automated Behavioural recording

After 96h exposure to the compounds, each 96-well plate was placed in ZebraLab to automatically record the locomotor activity of larvae with the help of VideoTrack software (both from View Point, S.A., Lyon, France). A light-emitting diode (LED) panel illuminated the 96-well plate from below. Recording was done under infrared light which, like the LED panel, is a fixed component of the ZebraLab system. The white light intensity of the ZebraBox was 500 lux. Locomotor activity was assessed by a subtraction method used for detection of objects darker than background with a minimum object size. A threshold of 0.1 mm (minimum distance moved) was used for filtering all of the data to remove system noise. Locomotor endpoints were designed to express the changes in the general swimming activity in response to light-dark stimulus.

A short test comprising of 14 minutes, called ‘visual motor response test’ was performed at 6 dpf as described elsewhere (55). All experiments were done at optimum temperature of 28 ± 0.5^◦^C. The visual motor response test has been previously used as frequently alternating periods of light and dark for a very short duration (not more than 10 minutes). This test is used to check abrupt change of locomotor activity (also called as visual startle response) after sudden shift from light to dark (38, 55-57). The experimental recording protocol consisted of three phases. First two minutes were given in the ZebraLab to acclimatize in the new environment. This phase was necessary to make sure that basal locomotor activity of zebrafish larvae is without any bias resulting in handling of the plate or change of location and hence was not used in the further analysis. After this acclimatization, the basal phase started, and consisted of 4 minutes to measure the basal locomotor activity while light in the ZebraLab remained ON. Immediately after basal phase, the lights were suddenly turned off for 4 min to record sudden change of locomotor activity which is called as ‘challenge phase’. Behavioural activity in the dark was also automatically recorded during this period with the help of infrared light. A third phase called ‘recovery phase’ was started immediately for 4 min after challenge phase to give zebrafish larvae time to recover from shock of darkness. All three phases consisted of 4-min to prevent habituation, and also to obtain more robust responses.

### Endpoint

Total distance moved for each minute during the 14 minute period was recorded. Average distance moved was calculated in all 3 phases i.e. basal, challenge and recovery.

### Statistical analysis

Statistical analyses were performed using GraphPad Prism version 5.04 for Windows, GraphPad Software, San Diego California USA, www.graphpad.com. One-way ANOVA was performed to analyse effect of various compounds on hatching rate and effect of compounds on locomotor activity. A Dunnett’s post hoc test was used to analyse multiple comparisons.

## Results

### Hatching percentage

The hatching percentage was monitored from 48-72 hpf which is the normal hatching period (54). We divided the effects of compounds on hatching into three categories after doing one-way ANOVA followed by Dunnett’s multiple comparison test: (i) compounds which have no significant effect on hatching, namely 2,4-Dichlorophenoxyacetic acid [F_(4,10)_=0.75, p=0.5801], MCPA [F_(3,8)_=1.0, p=0.4411], barium chloride [F_(5,12)_=0.8, p=0.5705], hexazinone [F_(5,12)_=0.84, p=0.5464] and strontium chloride [F_(5,12)_=2.4, p=0.0994] (Figure 1); (ii) compounds which delayed hatching, namely dimethoate [F_(5,12)_=1029, p<0.0001], benzophenone [F_(2,6)_=422.3, p<0.0001], triclopyr [F_(2,6)_=558.3, p<0.0001], pendimethalin [F_(5,12)_=406.2, p<0.0001], mercuric chloride [F_(2,6)_=484, p<0.0001], stannic chloride [F_(5,12)_=795, p<0.0001], maneb [F_(2,6)_=993.5, p<0.0001], hydroquinone [F_(3,8)_=400, p<0.0001], acephate [F_(4,10)_=527.2, p<0.0001], gallium chloride [F_(2,6)_=2257, p<0.0001], erbium chloride [F_(5,12)_=253.2, p<0.0001], diazinon [F_(4,10)_=475, p<0.0001], molinate [F_(5,12)_=417.3, p<0.0001], zinc chloride [F_(5,12)_=950.7, p<0.0001] and bromacil [F_(4,10)_=975.8, p<0.0001] (Figure 2); (iii) compounds which accelerated hatching, namely methomyl [F_(6,14)_=484, p<0.0001], glyphosate [F_(2,6)_=206, p<0.0001], paraquat [F_(4,10)_=18.79, p<0.0001], and amitrol [F_(6,14)_=205.9, p<0.0001] (Figure 3).

**Figure 1.**
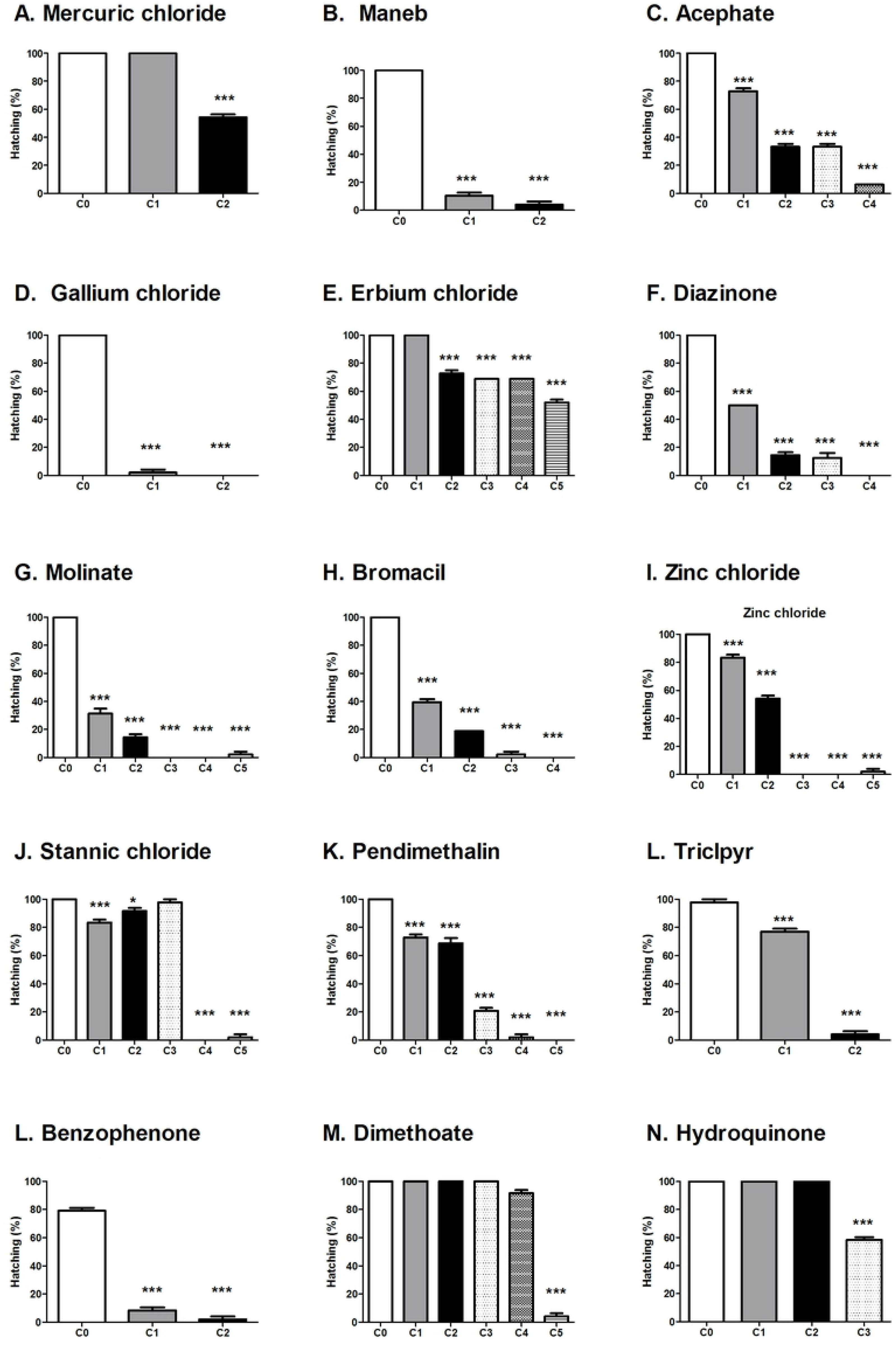
Hatching percentage after exposure to compounds that caused dose-dependent delay in hatching (as indicated by percent survivors hatched at 72hpf).

**Figure 2:**
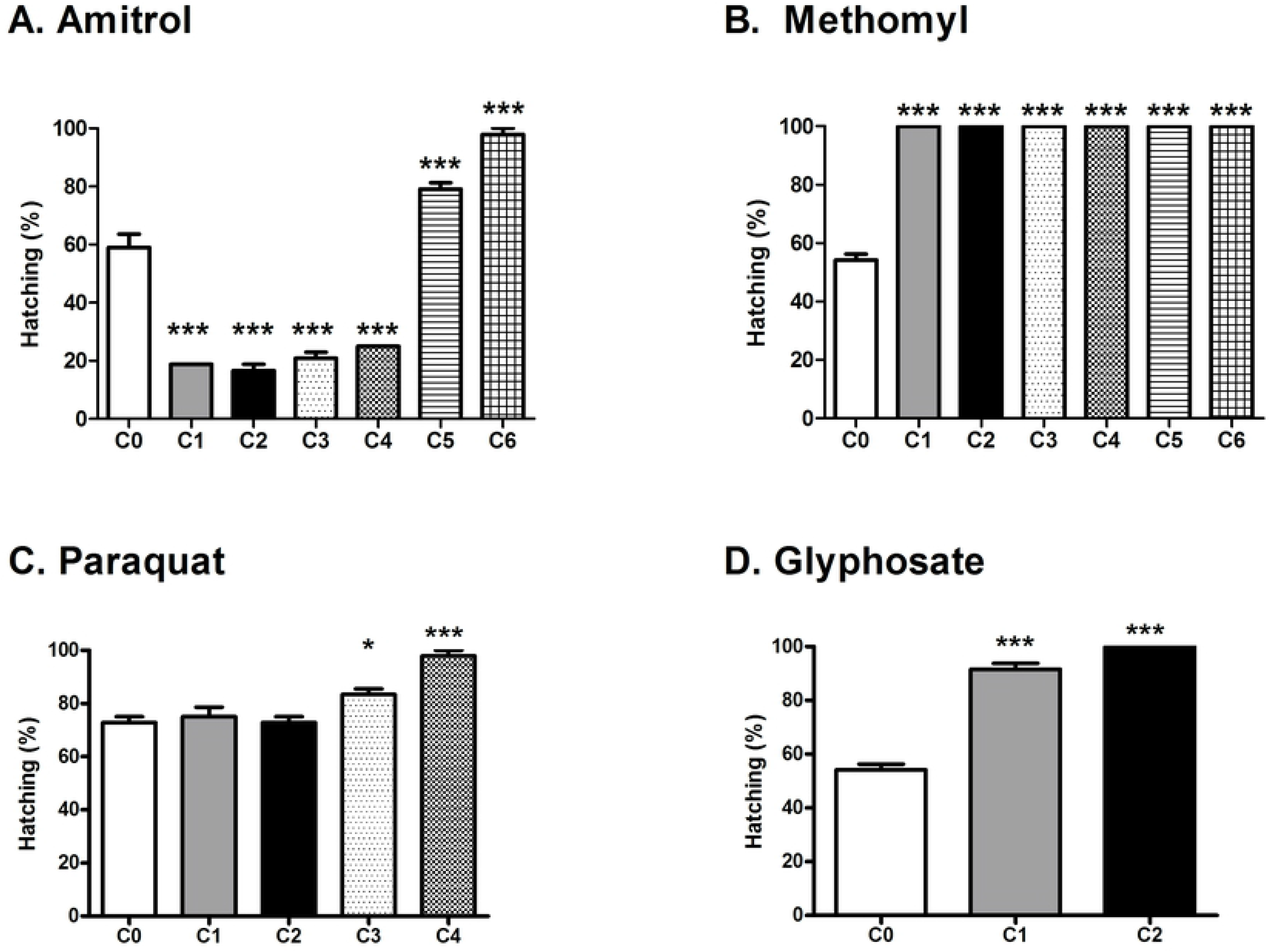
Hatching percentage after exposure to compounds that caused dose-dependent acceleration of hatching (as indicated by percent survivors hatched at 48hpf)

**Figure 3.**
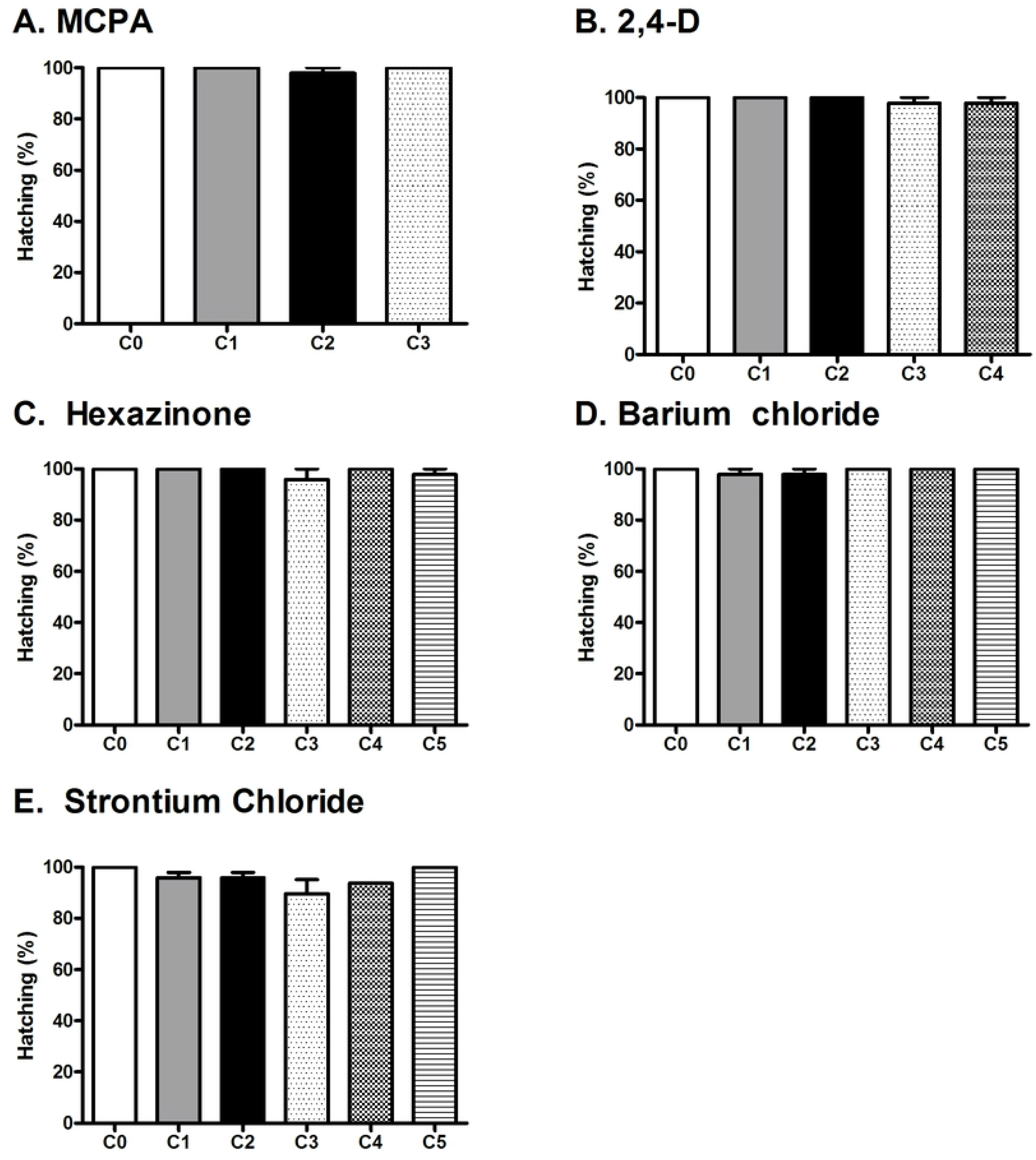
Hatching percentage after exposure to compounds that had no effect on hatching at 72hpf.

### Malformations

The malformations produced by the test compounds are summarized in Table 4. The features which were examined are yolk sac oedema, pericardial oedema, bent body, total length of the zebrafish larvae and pigmentation over the body. The compounds producing malformations in survivors were: Glyphosate, 2,4-Dichlorophenoxyacetic acid, diazinon, paraquat, methomyl and molinate (Table 4.) Mercuric chloride, gallium chloride and benzophenone produced lethality at all concentrations tested and hence malformations in survivors were not observed. The remaining compounds did not produce any of the malformations described in Table 1.

**Table 4:**
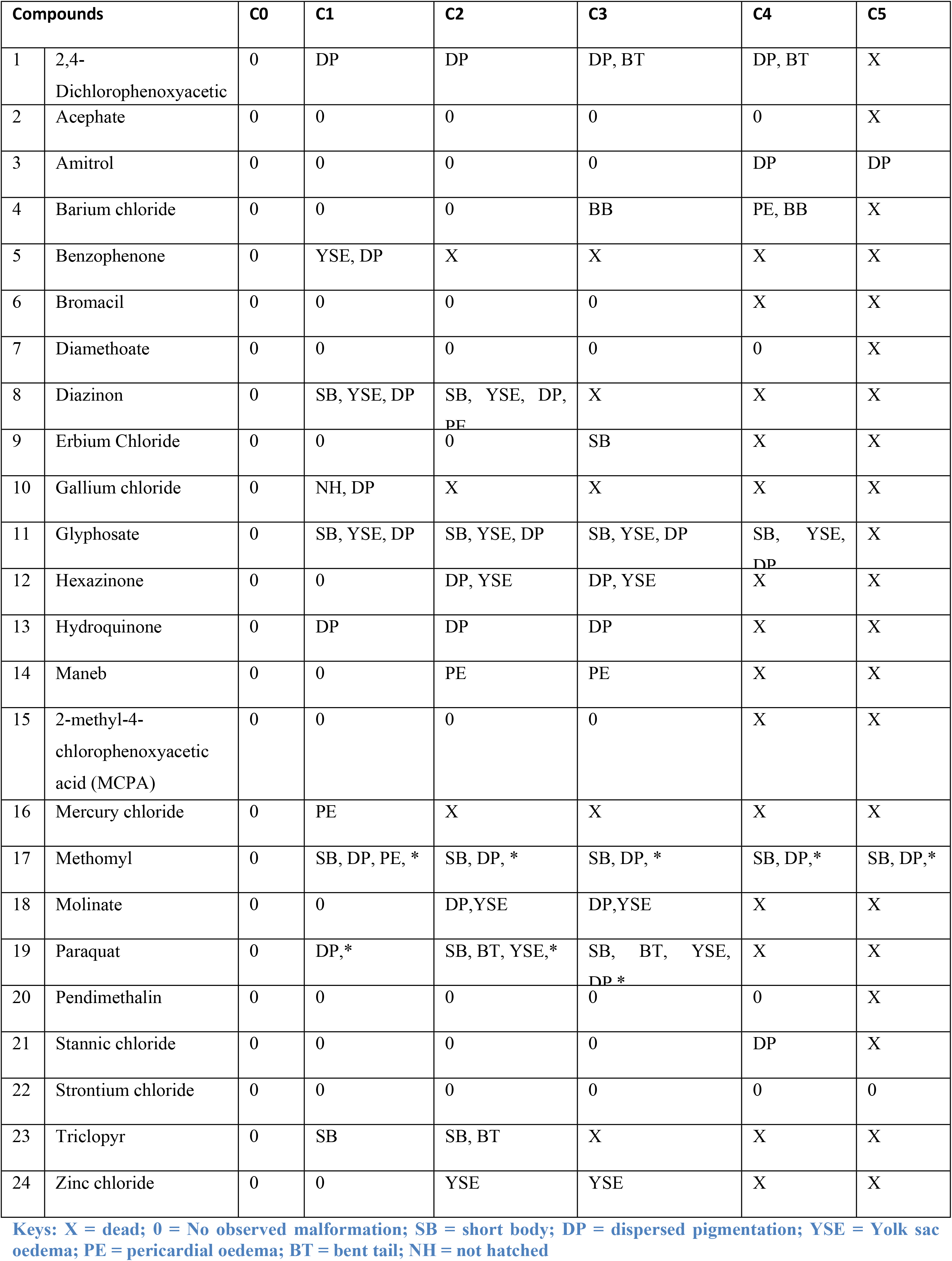
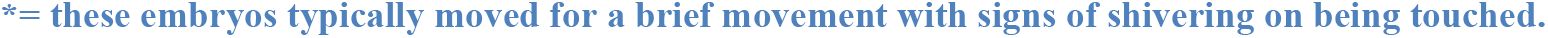
Malformations produced by varying concentrations (geometric series) of test compounds.

### LC_50_ value calculation and correlation with LD_50_ values of rodents

The LC_50_ values of zebrafish larvae determined after 96 h exposure to test compounds, and their corresponding LD_50_ values in rodents taken from the literature, are shown in Table 5. No correlation was found between the zebrafish and rodent values. Thus, a correlation test produced spearman’s rank correlation of −0.08498 (p=0.6999) and Pearson’s correlation −0.1086 (p=0.6218) between zebrafish embryo LC_50_ and rodent LD_50_ values.

**Table 5:** Zebrafish embryo LC_50_ values calculated in present study, and the corresponding rodent LD_50_ oral values based on the literature

| Compounds |  | Zebrafish embryo LC<sub>50</sub> (mg/L ±SEM) | Zebrafish embryo LC<sub>50</sub> (mmol/L) | Rodent LD<sub>50</sub> (mg/kg) | Rodent LD<sub>50</sub> (mmol/kg) |
| --- | --- | --- | --- | --- | --- |
| 1 | 2,4-Dichlorophenoxyacetic | 65.3±0.41 | 0.29±0.030 | 370(#) | 1.67 |
| 2 | Acephate | 5489.31±16.93 | 29.97±0.09 | 233(*) | 1.27 |
| 3 | Amitrol | ND | ND | 11000(*) | 130.83 |
| 4 | Barium Chloride | 135.35±3.26 | 0.65±0.03 | 132(*) | 0.63 |
| 5 | Benzophenone | 89.28±18.74 | 0.49±0.12 | 2895(*) | 15.89 |
| 6 | Bromacil | 270.2±0.67 | 1.03±0.071 | 5175(*) | 19.82 |
| 7 | Diamethoate | 684.3±2.04 | 2.97±0.002 | 60(*) | 0.26 |
| 8 | Diazinon | 27.5±0.38 | 0.09±0.024 | 96(*) | 0.32 |
| 9 | Erbium Chloride | 101.24±0.66 | 0.37±0 | 4417(Φ) | 16.14 |
| 10 | Gallium Chloride | 334.55±1.17 | 1.90±0.01 | 4700(*) | 26.69 |
| 11 | Glyphosate | 95.9±0.23 | 0.56±0.008 | 1568(*) | 9.27 |
| 12 | Hdroquinone | 4.40±0.33 | 0.04±0 | 245(*) | 2.23 |
| 13 | Hexazinone | 361.8±0.57 | 1.43±0.013 | 1690(*) | 6.7 |
| 14 | Maneb | 42.45±0.35 | 0.16±0 | 2600(*) | 9.80 |
| 15 | 2-methyl-4-chlorophenoxyacetic acid (MCPA) | 47.2±0.3 | 0.23±0.010 | 550(*) | 2.74 |
| 16 | Mercuric chloride | 0.27±0.00 | 0.001±0 | 6(⌘) | 0.02 |
| 17 | Methomyl | 59.7±0.39 | 0.367±0.030 | 10(*) | 0.06 |
| 18 | Molinate | 49.2±0.43 | 0.26±0.016 | 530(*) | 2.83 |
| 19 | Paraquat | 500.8±0.70 | 1.94±0.034 | 120(*) | 0.47 |
| 20 | Pendimethalin | 376.96±1.06 | 1.34±0.01 | 1340(*) | 4.76 |
| 21 | Stannic Chloride | 25841.98±2.30 | 0.81±0.01 | 46(*) | 0.18 |
| 22 | Strontium Chloride | 211.02±28.57 | 163.01±0.26 | 1874(☼) | 11.82 |
| 23 | Triclopyr | 69.24±0.17 | 0.27±0 | 729(*) | 2.84 |
| 24 | Zinc Chloride | 89.96±2.02 | 0.66±0.02 | 350(*) | 2.57 |

ND= Not determined
(\*) = from Hazardous Substances Data Bank at http://toxnet.nlm.nih.gov
(#) = from Extension Toxicology Network at http://extoxnet.orst.edu
(♣) = from ChemIDplus Advanced at http://chem.sis.nlm.nih.gov/chemidplus/cas/10138-41-7
(⌘) = from Material Safety Data Sheet at http://avogadro.chem.iastate.edu/MSDS/HgCl2.htm
(☼) = from http://www.guidechem.com/msds/10025-70-4.html

The relative toxicity ([zebrafish LC_50_ mmol/L] ÷ [rodent LD_50_ mmol/kg]) of individual compounds is shown in Figure 4. Compounds which were less toxic in zebrafish than in rodents inlcude bromacil, dimethoate, diazinon, glyphosate, haxezinone, MCPA, molinate, 2,4-Dichlorophenoxyacetic acid, acephate, barium chloride, benzophenone, erbium chloride, gallium chloride, hydroquinone, maneb, mercuric chloride, pendimethalin, triclopyr and zinc chloride. The compounds which were more toxic in zebrafish than in rodents were methomyl, paraquat, strontium chloride and stannic chloride.

**Figure 4:**
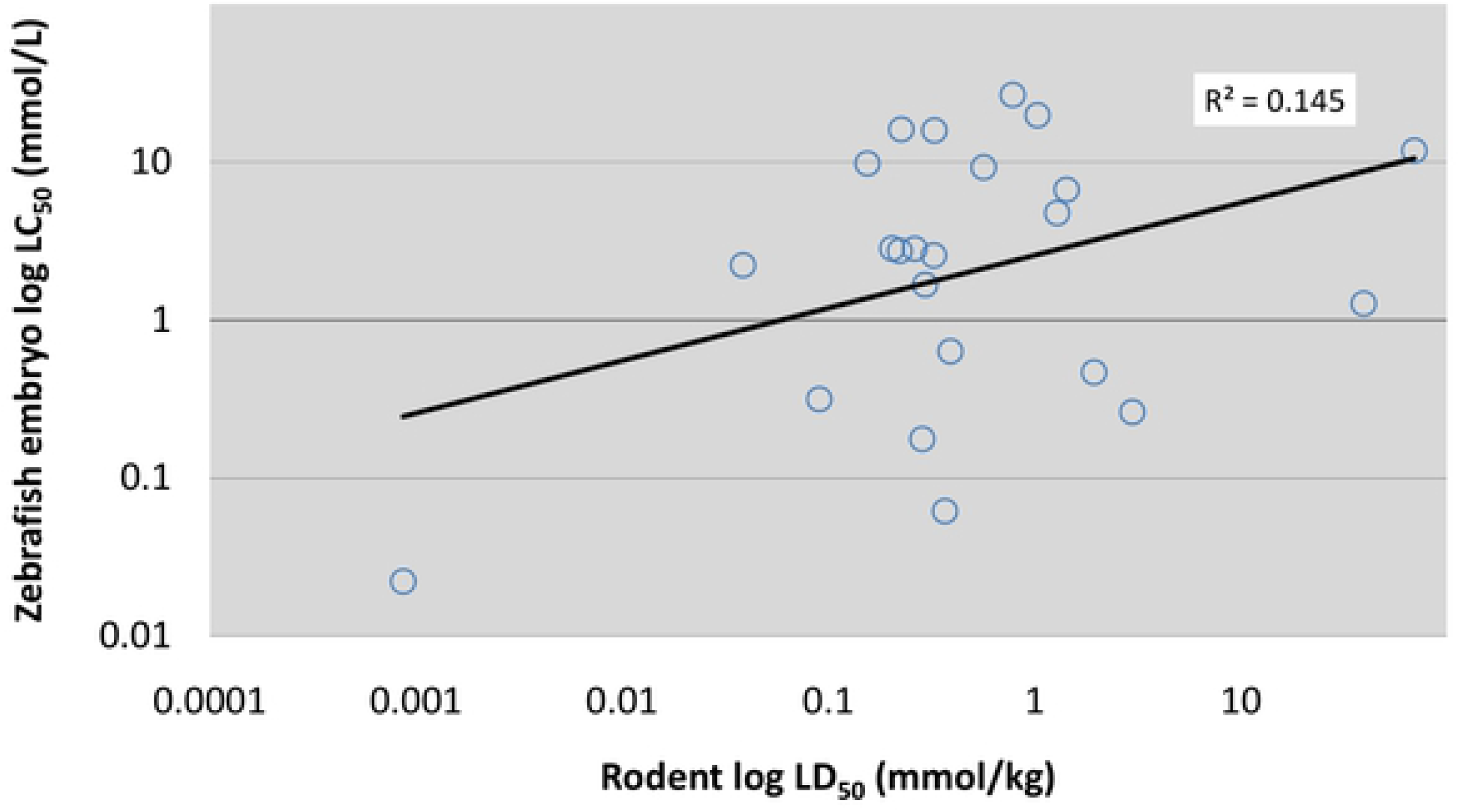
Relative toxicity of individual compounds tested in this study. Zebrafish embryo LC_50_ was determined based on cumulative mortality after 96 h exposure of compounds from three independent experiments and rodent LD50 was taked from the literature.

### Locomotor activity

The visual motor response test was used to assess the integrity of the central and peripheral nervous system together with visual and musculoskeletal system development. On the basis of the visual motor response test, four distinct responses were found as follows:

### Monotonic stimulation response

One-way ANOVA followed by Dunnet’s test for multiple comparisons showed that the locomotor activity of zebrafish larvae in the challenge phase was significantly increased (Figure 5). The compounds which showed monotonic stimulation response were: paraquat [F_(3,58)_=7.439, p<0.001], stannic chloride [F_(3,58)_=4.981, p=0.0038] and amitrol [F_(5,89)_=4.155, p<0.001].

**Figure 5:**
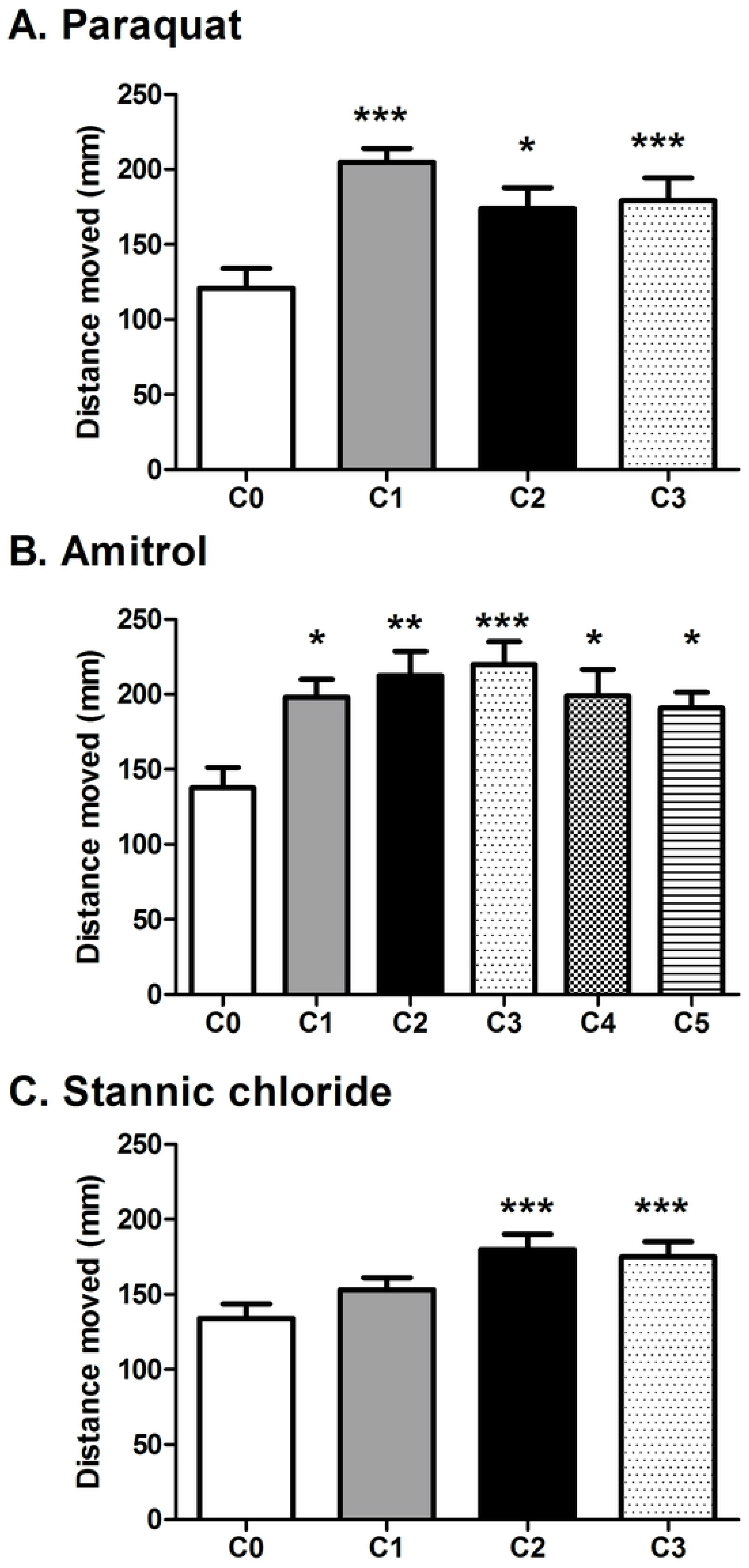
Distance moved during the challenge phase of the visual motor response test by zebrafish larvae at 5dpf. All these compounds displayed a significant concentration-dependent increase in distance moved. Error bars represent ±SEM of N=48 control and survived embryos for each concentration of each compound from three independent experiments. Statistical icons: *=p<0.05, **=p<0.01 and ***=p<0.001

### Monotonic suppression response

One-way ANOVA test followed by Dunnet’s post hoc test for multiple comparisons showed that an increase in the concentration of the compounds caused suppression of locomotor activity (Figure 6). The locomotor activity of zebrafish larvae was significantly decreased in the challenge phase as compared to controls. The compounds with monotonic supression were: strontium chloride [F_(5,78)_=37.90, p<0.0001], zinc chloride [F_(3,53)_=7.506, p<0.001], pendimethaline [F_(3,48)_=28.13, p<0.0001], diazinon [F_(2,42)_=5.267, p<0.001], hexazinone [F_(3,49)_=16.08, p<0.0001], methomyl [F_(3,27)_=25.75, p<0.0001], molinate [F_(3,51)_=20.61, p<0.0001], dimethoate [F_(3,60)_=13.31, p<0.0001] and barium chloride [F_(4,63)_=10.80, p<0.0001].

**Figure 6.**
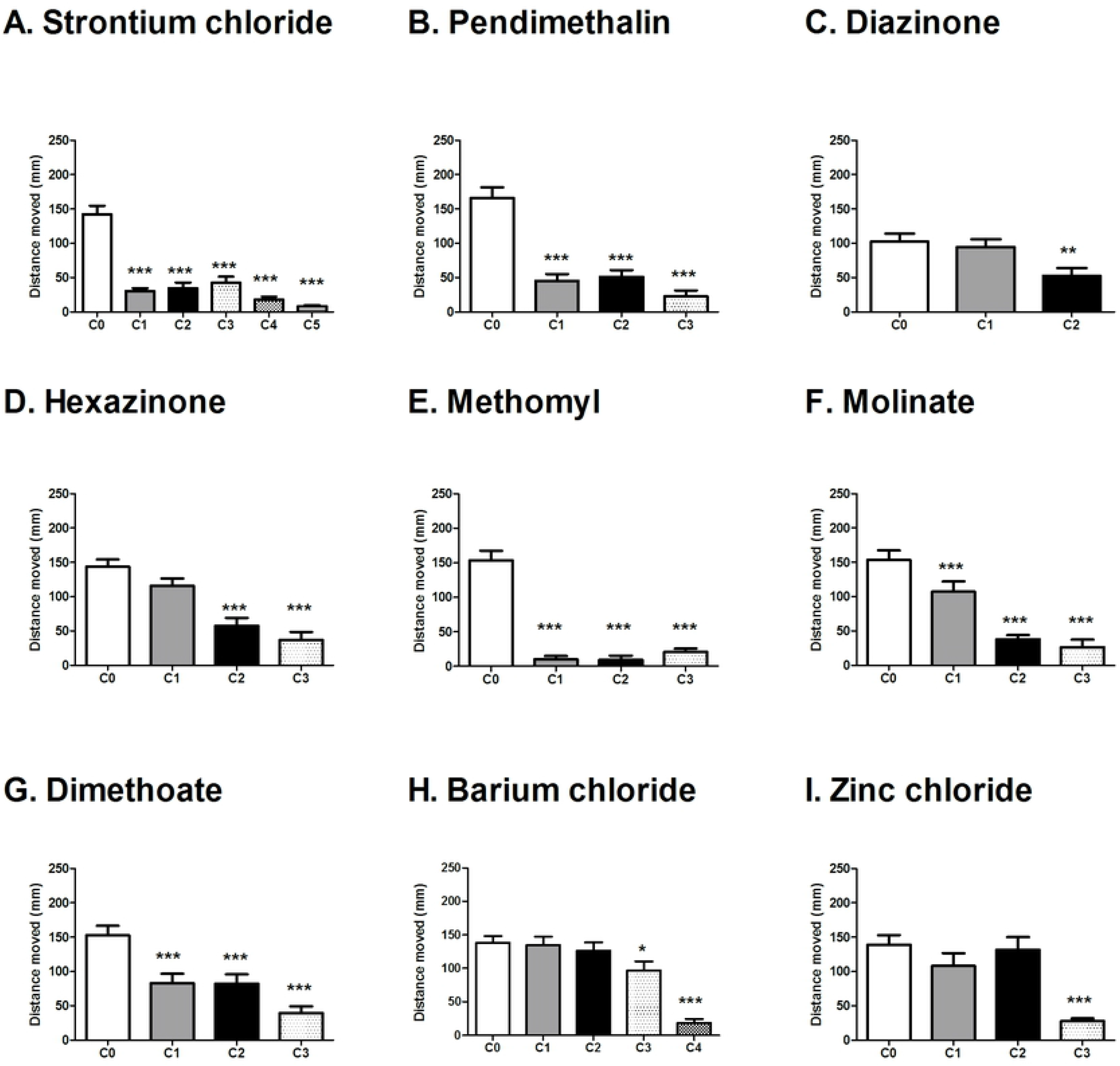
Distance moved during the challenge phase of the visual motor response test by zebrafish larvae at 5dpf. These compounds showed a significant concentration-dependent decrease in the locomotor activity. Error bars represent ±SEM of N=48 control and survived embryos for each concentration of each compound from three independent experiments. Statistical icons: *=p<0.05, **=p<0.01 and ***=p<0.001

### Biphasic response (dose dependent stimulation and suppression)

One-way ANOVA test followed by Dunnet’s post hoc test for multiple comparisons showed a significant difference in the locomotor activity of some compounds, at certain concentrations tested, and controls (Figure 7). In these cases, the locomotor activity increased with increasing concentration, and then decreased at yet higher concentrations. The compounds with this biphasic response were erbium chloride [F_(3,58)_=20.28, p<0.0001], 2,4-Dichlorophenoxyacetic acid [F_(4,66)_=9.143, p<0.0001] and hydroquinone [F_(3,56)_=12.58, p<0.0001].

**Figure 7:**
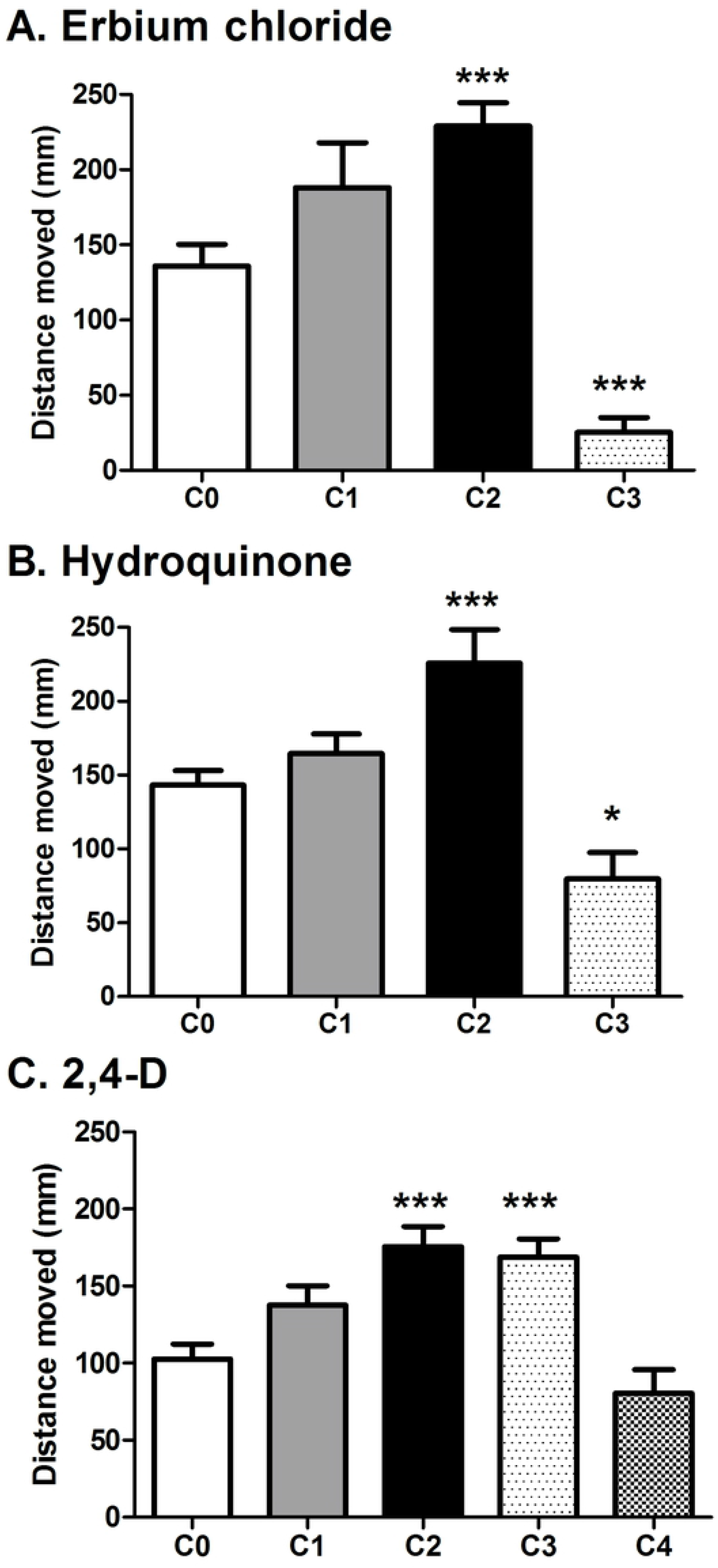
Distance moved during the challenge phase of the visual motor response test by zebrafish larvae at 5dpf. These compounds showed a significant concentration-dependent increase and then a decrease at a high concentration in the locomotor activity. Error bars represent ±SEM of N=48 control and survived embryos for each concentration of each compound from three independent experiments. Statistical icons: *=p<0.05, **=p<0.01 and ***=p<0.001

### No effect

For some compounds, the locomotor activity of zebrafish larvae was unaffected, regardless of concentration tested (Figure 8). One-way ANOVA test followed by Dunnet’s post hoc test for multiple comparisons showed no significant difference in the locomotor activity between the various concentrations of compounds. The compounds without any effect on the locomotor activity were: maneb [F_(3,55)_=2.26, p=0.0908], glyphosate [F_(4,73)_=0.5964, p=0.6664], MCPA [F_(3,59)_=2.272, p=0.0895] and bromacil [F_(2,42)_=2.154, p=0.1287].

**Figure 8.**
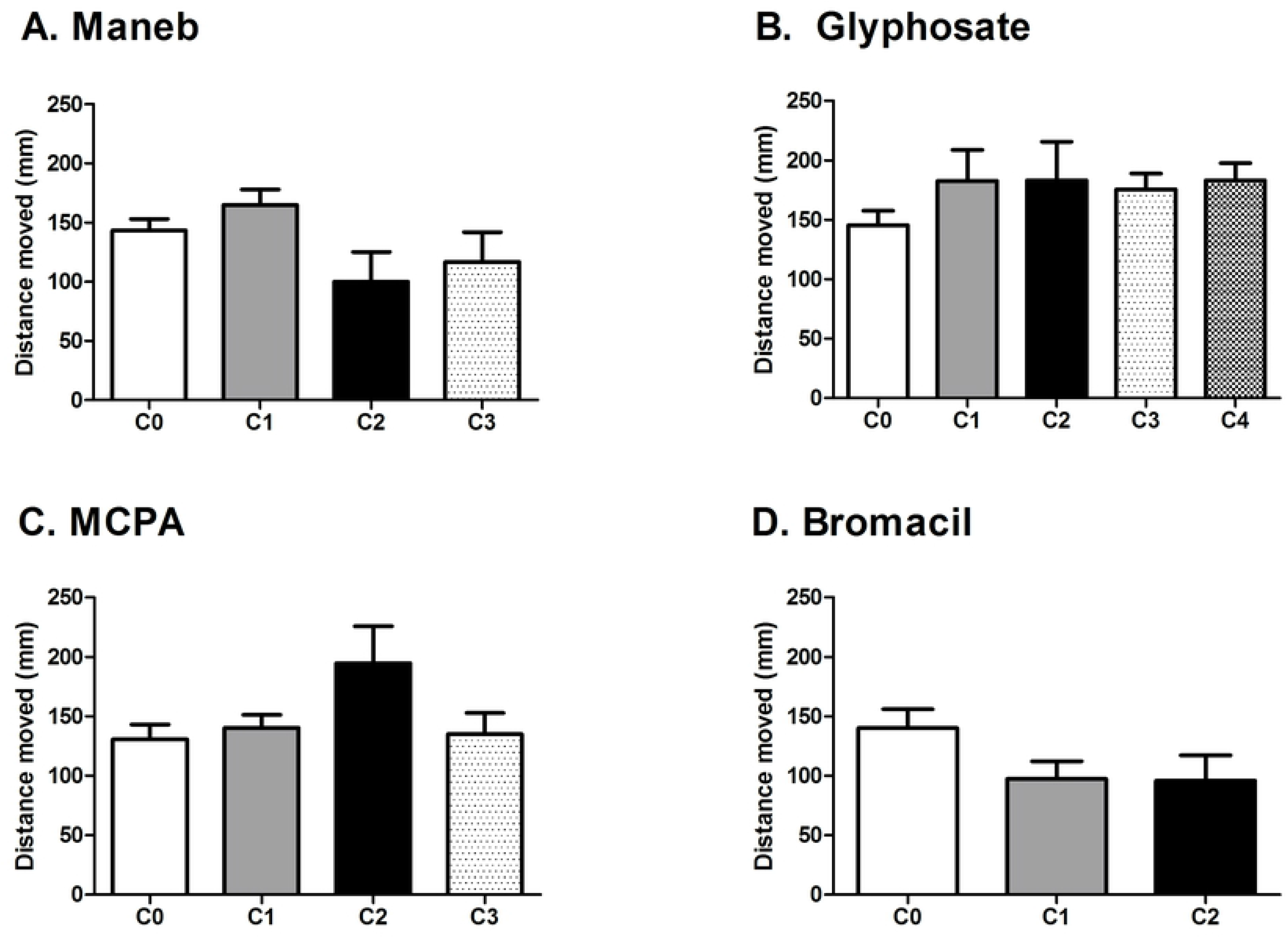
Distance moved during the challenge phase of the visual motor response test by zebrafish larvae at 5dpf. These compounds showed no significant difference in locomotor response as compared to control. Error bars represent ±SEM of N=48 control and survived embryos for each concentration of each compound from three independent experiments. Statistical icons: *=p<0.05, **=p<0.01 and ***=p<0.001

## Discussion

### Hatching

The first significant finding in the present study is that the differential hatching percentage depends on the compound tested. Hatching is an essential step in zebrafish development, and delayed hatching makes zebrafish more susceptible to predators; complete inhibition of hatching may also result in death (58). We found that the time of hatching is influenced by compound type, and by concentration. Many compounds tested resulted in delayed hatching (compared to controls). However, four compounds were associated with accelerated hatching, namely: amitrol, methomyl, paraquat and glyphosate. With amitrol, lower concentrations delayed hatching, while higher concentrations accelerated it. By contrast, lower concentration of paraquat did not have effect on hatching while higher concentrations accelerated the hatching as compared to control larvae. The higher concentrations of methomyl and glyphosate also accelerated the hatching. Fourteen compounds out of the 24 tested had no significant effect on the hatching rate.

Hatching in zebrafish takes place in two steps. The first step is the release of hatching enzyme by the hatching gland which breaks down the inner vitelline envelope of the acellular chorion (59). The seconds step is the spontaneous movement of the embryo which starts around 19hpf until the hatching. The delayed hatching in the present case might be due to delay in the release of hatching enzyme or a delay in the spontaneous movement activity. The other explanation lies in the presence of chorion around the zebrafish embryo. The 3.5 µm thick chorion (60) protects the zebrafish embryo against the toxic effects of compounds (61), and acclimation of different toxins (62). It is even possible that delayed hatching might allow the embryo to survive short-term exposure of compounds, which would have killed the hatched (non-chorion-protected) larvae.

It remains to be elucidated how these chemicals can accelerate or inhibit the hatching process, and what the ecological consequences might be in the wild. However, this phenomenon shows that the embryo can react to chemicals at concentrations at which larval survival is not affected. Although the mechanism and consequences of delayed or accelerated hatching are unknown, it is possible that hatching time may serve as a sublethal response variable for embryonic development in toxicity tests. Further work is required to examine these issues.

### Morphological malformations

It has been found that the physical properties of chemicals did not fully predict lethality or developmental outcomes; rather, individual outcomes such as pericardial oedema and yolk sac oedema are more reliable indicators of developmental toxicity (22). Thus, in order to see the teratogenic effects of compounds, we screened for malformations. It was found that 29% (7/24) of the compounds produced none of the morphological abnormalities in the zebrafish embryos described in Table 1 at any concentration. By contrast, 71% (17/24) compounds produced various malformations summarized in Table 4. The most common abnormality in these larvae was dispersed pigmentation on the body; this is considered an indication of stress (63). The compounds 2,4-Dichlorophenoxyacetic acid, paraquat and barium chloride produced axial curvature and deformed or bent tail. It has been suggested that these types of malformation might be due to delayed hatching (64), a conclusion consistent with the results of the present study.

Acephate, amitrol, strontium chloride, stannic chloride, bromacil, dimethoate, MCPA and erbium chloride caused no morphological deformities at any concentration. By contrast, diazinon, glyphosate, hexazinone, methomyl, molinate, 2,4-Dichlorophenoxyacetic acid were among the most teratogenic compounds tested resulting in multiple malformations. Benzophenone, gallium chloride and mercuric chloride simply produced lethality.

### LC_50_ of zebrafish vs. LD_50_ of rodents

In the present study, the correlation between LC_50_ of zebrafish and LD_50_ of rodents was very weak for metals and biocides considered together (R2=0.1456). We compared the LC_50_ of zebrafish with oral LD_50_ in rodents taken from the literature. Where data were available from more than rodent species, we did not take the average, but used a single value from one study.

The difference we find between LC_50_ of compounds in zebrafish and oral LD_50_ in rodents can be explained by various factors. The first factor is that we are comparing the developmental toxicity of a compound in the zebrafish embryo versus a rodent adult. Thus we are comparing different life stages. Secondly, the route of exposure should also be taken to into account. In case of the zebrafish embryos, we exposed chronically to compound for 96 h beginning at 24hpf. In the early part of this period, there is a relatively impermeable chorion (3.5 µm thick, composed of three acellular layers) surrounding the embryo (60). After hatching, the drug could, in principle, be absorbed through the skin, taken up by the gills, or absorbed from the pharynx or gut. Little is known about the absorption of drugs by zebrafish embryos. In the case of the rodent studies used here for comparison, compounds were administered orally. An important issue for futures studies using the zebrafish embryo model is to examine the route of absorption of compounds from the environment and to compare it with absorption in rodents and other mammals from the digestive tract or other routes.

It has been reported that zebrafish LC_50_ values of a variety of compounds correlate well with the corresponding LD_50_ values in rodents (9, 65) and birds (66). On this basis, it has been suggested that zebrafish embryos/larvae are a good alternative method for developmental toxicity studies (67). However, it has also been emphasised that special care should be taken in considering predictivity because this parameter varies with the class of compounds (9). The authors showed that the slope of the regression line (zebrafish LC_50_ vs. rodent LD_50_) varied from 0.36 to 1.27 depending on the compound class. In another study, Parng and colleagues (65) showed that LC_50_ values of 11 out of 18 compounds were correlated with the LD_50_ values of those compounds in mice. Together, these studies suggest that the predictivity of the zebrafish embryo model is critically dependent on compound class.

Another example of comparative toxicology in the zebrafish model (68) used multiple approaches to study cell cycle inhibition of various compounds. Zebrafish embryos were tested to screen 16,320 compounds to assess the level of serine-10-phosphorylated-histone 3. They also tested 17 known chemicals which can disrupt the cell cycle in mammals, and found that 9 out of 17 compounds were positive. The other 8 chemicals were active in the *in vitro* AB9 zebrafish fibroblast culture preparation making a total of 94% of tested compounds that were active in zebrafish assays. Thus, the authors concluded (68) that the drug target conservation between zebrafish and mammals is very high.

In summary, our results, together with other studies, suggest that although the zebrafish embryo is a valid alternative/complimentary model in toxicity studies, its use as a surrogate to predict rodent and human acute toxicity can depend strongly on the compound type.

### Locomotor activity

In order to see the effect of compound type on locomotor activity, we used the visual motor response test at 5dpf. This test has previously proved effective as a simple locomotor behaviour test for assessing effects of compounds. (38, 40, 42, 55, 69). We chose larvae at 5dpf, a time point at which they display a wide range of behavioural repertoires, and at which many organs are differentiated (70).

A number of compounds that we tested showed a significant concentration-dependent suppression of locomotor activity in the visual motor response test. These include agents that have a comparable effect in rodents. Pendimethalin and methomyl supressed the locomotor activity in the zebrafish larvae in the present study, and also in rodents (71, 72). The effect of a few compounds on the locomotor activity of rodents and zebrafish larvae is shown in Table 6.

**Table 6:** Comparison of effects of selected compounds on zebrafish and rodents locomotor activity. The effect on zebrafish larvae are derived from the present study while the effect on rodent is derived from the literature.

| Compound | Effect of compound on zebrafish locomotor activity in present study | Effect of compound on rodent locomotor activity (from literature) | References |
| --- | --- | --- | --- |
| Pendimethalin | Decreased | Decreased | (71) |
| Methomyl | Decreased | Decreased | (72) |
| Dimethoate | Decreased | Decreased | (80) |
| Hydroquinone | Decreased | Decreased | (77) |
| 2,4-Dichlorophenoxyacetic | Decreased | Decreased | (78) |
| Paraquat | Increased | Increased | (81) |
| Maneb | No effect | Decreased | (82) |

On the other hand, some compounds increased the locomotor activity in the challenge phase as compared to controls in our study. Specifically, zebrafish larvae treated with amitrol, stannic chloride and paraquat showed hyperactivity in a concentration dependent manner. Paraquat-induced toxicity has been linked to Parkinson’s-like neurological degenerative mechanisms both in rats (73) and in zebrafish (74). It is possible that the hyperactivity of zebrafish larvae recorded in this study in the challenge phase was due to Parkinson-like tremors. Further work is required to examine this possibility.

Some compounds in this study showed a biphasic effect, that is, either stimulation or suppression of locomotor activity depending on the concentration. For example, erbium chloride, hydroquinone and 2,4-dichlorophenoxyacetic acid increased the locomotor activity in a concentration dependent manner at lower concentrations, but suppressed it at higher concentrations. A biphasic response has also been observed in rodents following exposure to toluene (75) and ethanol (76). Hydroquinone in rodents has been known to decrease locomotor activity (77). Similarly, 2,4-dichlorophenoxyacetic acid is also known to decrease the spontaneous locomotor activity in rats, contrary to our results where it was increased initially before decreasing at higher dose (78). Possible explanations for the different responses in zebrafish in this study, compared to the rodent literature, could include the different route of exposure, as well as different concentrations in the tissues. Again, these findings emphasise the need for comparative studies of absorption of compounds in the zebrafish embryo.

When exposing zebrafish embryos to toxicants, there are several possible mechanisms for the effect on locomotor behaviour. For example, the toxicant could cause retarded development of the locomotor and nervous systems, and the latter could include visual impairment. Visual impairment has been implicated in the effects of ethanol on zebrafish because it causes abnormalities of eye development (i.e. microphthalmia; see (55).

Hypoactivity can also be attributed to other malformations (79). However, the presence of malformations cannot explain the hypoactivity seen in the present study after treatment with pendimethalin, strontium chloride and dimethoate, in which no malformations were present. In contrast, we found that larvae exposed to glyphosate were severely malformed but showed no difference in locomotor activity. In conclusion, there are multiple factors which can contribute to the hyper-or hypoactivity in the zebrafish larvae and a single factor cannot explain all the variations in locomotion.

## Conclusion

We have shown that different classes and even different compounds within the same class produce a range of different effects on zebrafish. Hatching was either delayed or accelerated depending on the compound, and the compounds produced varying malformations during development at difference concentrations. Zebrafish larvae showed three types of behavioural responses: (i) hypoactivity; (ii) hyperactivity; and (iii) biphasic response (a dose-dependent shift between hypo- and hyperactivity). When LC_50_ values of compounds were compared to published LD_50_ values in rodents, they showed poor correlation. It can be suggested that although the zebrafish embryo model has been embraced by wide scientific community as an alternative model for screening the developmental toxicity potential of compound, its predictivity for mammalian toxicity needs to be determined per compound class. More work is required to draw a general conclusion about predictive power of zebrafish model.

